# CRISPR delivery particles for developing therapeutic strategies in metabolic disease

**DOI:** 10.1101/352799

**Authors:** Yuefei Shen, Jessica L. Cohen, Sarah M. Nicoloro, Mark Kelly, Batuhan Yenilmez, Felipe Henriques, Emmanouela Tsagkaraki, Yvonne J.K. Edwards, Xiaodi Hu, Randall H. Friedline, Jason K. Kim, Michael P. Czech

## Abstract

RNA-guided engineered nucleases derived from a prokaryotic adaptive immune system known as CRISPR-Cas represent a promising platform for gene deletion and editing. As a therapeutic approach, direct delivery of Cas9 protein and guide RNA could circumvent the safety problems associated with plasmid delivery and therefore represents an attractive tool for genome engineering. Gene deletion or editing in adipose tissue to enhance its energy expenditure, fat oxidation and secretion of bioactive factors through a “browning” process presents a potential therapeutic strategy to alleviate metabolic disease. Here, we developed novel CRISPR delivery particles, denoted CriPs, composed of nano-size complexes of Cas9 protein and single guide (sg)RNA, coated with an amphipathic peptide called Endo-Porter that mediates entry into cells. Efficient CRISPR-Cas9 mediated gene deletion of ectopically expressed Green fluorescent protein (GFP) by CriPs was achieved in multiple cell types including a macrophage cell line, primary macrophages and primary pre-adipocytes. Significant GFP loss was also observed in peritoneal exudate cells with minimum systemic toxicity in GFP expressing mice following intraperitoneal injection of CriPs containing sgRNA targeting *Gfp*. Furthermore, the disruption of the *Nrip1* gene in white adipocytes by CriPs enhanced adipocyte “browning” with a marked increase of UCP1 expression. Deletion of *Nrip1* by CriPs did not produce detectable off-target effects. Thus CriPs represent a novel CRISPR delivery system for Cas9 and sgRNA that is effective for ablating targeted gene products in cultured cells and *in vivo,* and provide a potential therapeutic strategy for metabolic disease.

## INTRODUCTION

Gene editing based on the clustered regularly interspaced short palindromic repeats (CRISPR) – associated protein 9 (Cas9) system presents significant therapeutic potential for treating a wide range of diseases(1–4). The CRISPR-Cas9 system, containing the RNA-guided nuclease (Cas9 protein) and a single guide RNA (sgRNA), recognizes a protospacer-adjacent motif (PAM) and generates double-stranded DNA breaks (DSBs) at 3 base pairs upstream of a PAM site(5). DSBs are repaired by non-homologous end joining (NHEJ) to generate permanent gene deletion by inducing random insertions and deletions (indels) and by homology directed repair (HDR) to correct gene mutations with the use of a guide template DNA(5). A key challenge for CRISPR-Cas9 based therapeutics is the efficient, safe delivery of genome editing macromolecules to achieve eventual translation to clinical efficacy and safety(6–9). Viral vectors including adeno-associated virus (AAV) have shown efficient *in vivo* delivery and expression of CRISPR-Cas9(10–13). However, it is difficult to fit coding sequences for Streptococcus pyogenes Cas9 (SpCas9) plus sgRNAs into AAV vectors due to the restricted packaging capacity of AAVs(14). AAV-based Cas9 delivery also tends to cause significant off-target genome damage due to the sustained expression of Cas9(15, 16). In addition, the immune response to AAV capsids and the immunogenicity of the long-term existing bacterial Cas9 protein can limit their applications in human(11). Physical delivery approaches of CRISPR-Cas9, such as electroporation(17–19), micro-injection(20) and hydrodynamic injection(21, 22), have also been successfully used, but with concerns such as cell viability, toxicity and difficulty to apply *in vivo*.

The limitations associated with both viral delivery and physical delivery can be addressed using non-viral delivery systems such as lipid nanoparticles(23–27), DNA nanoclew(28), gold nanoparticles(29, 30), as well as chemically conjugating Cas9 protein with polymers(31) and cell penetrating peptides(32). Furthermore, direct delivery of Cas9-sgRNA ribonucleoprotein (RNP) is being considered as a promising therapeutic strategy. Cas9-sgRNA RNP delivery could circumvent the safety problems associated with plasmid delivery such as uncontrolled integration of DNA segments into the host genome and unwanted immune response to plasmids encoding Cas9 protein and sgRNA(6). Therefore, non-viral delivery of Cas9-sgRNA RNP represents an attractive tool for genome engineering. Cas9-sgRNA RNP delivered by non-viral delivery systems has been tested in cell culture(30, 33), primary cells(15, 19) and for local delivery such as inner ear injection(4, 23), skin injection(18), intra-tumor injection(28), intramuscular injection(29) and intracranial injection(34). However, to the best of our knowledge, Cas9-sgRNA RNP has not yet been tested systemically *in vivo* using a fully non-viral delivery system.

Application of CRISPR in therapies for type 2 diabetes would be attractive since this malady and its complications afflicts around 30 million adults in the United States and is a leading cause of death(35). White adipose tissue (WAT) stores triglycerides and expands greatly during the onset of obesity, which can prompt insulin resistance, failure of insulin secretion and the development of type 2 diabetes(36). Unlike WAT, brown adipose tissue (BAT) is composed of brown adipocytes that display high capacity for fat oxidation and a high number of mitochondria containing uncoupling protein 1 (UCP1) for nonshivering thermogenesis that plays a beneficial role in metabolism(37). BAT can also secrete beneficial factors to increase glucose uptake and fatty acid oxidation in other tissues(38, 39). Recent data indicate that increased BAT can favorably control whole-body glucose homeostasis and is associated with lean, insulin sensitive phenotypes(40–42). White adipocytes can be converted to brown or “beige” adipocytes by silencing molecular targets that suppress energy expenditure, fatty acid oxidation and insulin signaling, such as the nuclear co-repressor *Nrip1* gene(43, 44) (also denoted as RIP140). *Nrip1* silencing by RNAi in white adipocytes leads to adipocyte “browning” and enhances fatty acid oxidation, mitochondrial respiration and insulin mediated glucose uptake(43). *Nrip1* null mice present lean phenotypes with improved insulin sensitivity and glucose tolerance(45), suggesting that *Nrip1* is a powerful molecular target for alleviating type 2 diabetes. However, NRIP1 is not easily “druggable” by small molecules. Side effects may occur if inhibition of *Nrip1* is not restricted to adipocytes and muscle(46). Thus, an effective therapeutic strategy for targeting *Nrip1* requires its gene deletion specifically in adipocytes or skeletal muscle or both, rather than in other tissues or through whole-body targeting.

Here, we developed a novel CRISPR delivery system, denoted CRISPR delivery particles (CriPs), composed of nano-size complexes of the CRISPR components Cas9 protein and sgRNA targeting a gene of interest, complexed with an Endo-Porter (EP) peptide that mediates entry into cells. EP is an amphipathic peptide composed of leucine and histidine residues, which we have previously used successfully for targeted siRNA delivery in cells(47–50). It is hypothesized that the weak-base histidine residues of EP facilitate the endosomal escape of the cargoes by the “proton-sponge effect”(51). As proof of concept, efficient CRISPR-Cas9 mediated gene deletion of the green fluorescent protein (*Gfp*) gene was observed in multiple cell types isolated from GFP transgenic mice including primary macrophages and primary pre-adipocytes. GFP loss was achieved in about 50% of macrophages and primary pre-adipocytes as determined by flow cytometry analysis, in response to treatment with CriPs targeting *Gfp*. Indels in the *Gfp* genomic locus were confirmed by measurements using a T7 endonuclease I (T7E1) assay. Significant GFP loss was also observed in peritoneal exudate cells (PECs) isolated from GFP transgenic mice after 5 daily intraperitoneal (i.p.) injections with CriPs targeting *Gfp* as determined by flow cytometry and confirmed by nucleotide sequencing. To our knowledge, this is the first study suggesting that Cas9-sgRNA RNP can be systemically delivered *in vivo* by a simple non-viral system to achieve significant gene deletion efficiency. Furthermore, deletion of the *Nrip1* gene in white adipocytes by CriPs converted them to a more “brown” adipocyte phenotype, with a remarkable increase of UCP1 expression and no detectable off-target effects as determined by T7E1 assay.

## RESULTS

### Design and Characterization of CriPs

The goal of the present study was to develop a simple delivery vehicle to effect specific gene deletion via the CRISPR-Cas9-based genome targeting mechanism. For therapeutic applications, we aimed to directly deliver Cas9 protein rather than plasmids that encode the protein to overcome such problems as uncontrolled integration of plasmid DNA into the host genome, unwanted immune responses and safety issues. Here, we report the preparation of CriPs that can deliver Cas9 protein bound to sgRNA to mediate gene deletion *in vitro* and *in vivo* (Fig. 1). Purified bacterial Cas9 protein that is carefully processed to remove endotoxin and other contaminants is used for loading of sgRNA. The sgRNA sequence is designed to target a selected gene at a site adjacent to a PAM sequence in the DNA of that target gene. The loaded Cas9-sgRNA complexes are then complexed with amphipathic EP peptide, which is designed to mediate the uptake of the Cas9-sgRNA complexes into live cells without toxicity or detectable damage. The inert CriPs contain only the three molecular components shown in Figure 1 (Cas9 protein, sgRNA and EP). As opposed to gene therapy approaches, no permanent viral vectors or genetic insertions of viral DNA are utilized. Cas9 protein that enters cells through transfection mechanisms lasts no longer than 24 hours within cells(52). The EP peptide and the sgRNA are also rapidly degraded within cells. Thus, for *ex vivo* therapeutic approaches such as cell therapy and transplantation of *ex vivo* engineered tissue, Cas9 and other components in CriPs are not present in cells that are implanted. Additionally, unlike RNAi-based approaches, CriPs-mediated gene editing lasts for the lifespan of a cell, which allows for less frequent therapeutic administration and less chances of immune response or chronic toxicity.

**Figure 1.**
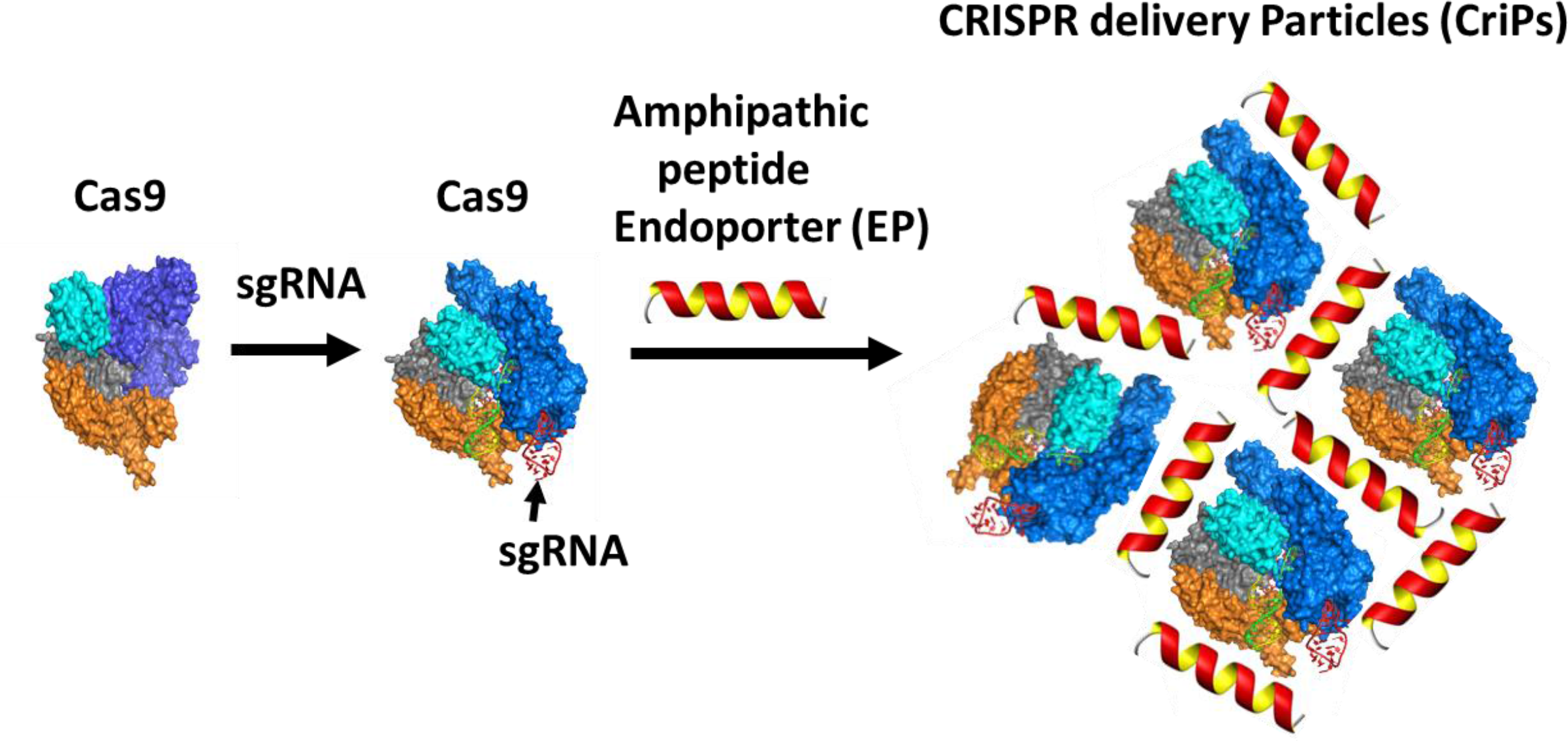
Preparation of CRISPR delivery particles (CriPs). Purified bacterial Cas9 protein that is carefully processed to remove endotoxin and other contaminants is used for loading of sgRNA. The sgRNA sequence is designed to target a selected gene at a site adjacent to a PAM sequence in the DNA of that target gene. The loaded Cas9-sgRNA nanocomplexes are then coated with an amphipathic peptide, denoted as Endo-Porter (EP), which is required to mediate uptake of the Cas9-sgRNA complexes into live cells without toxicity or detectable damage.

CriPs with different ratios of EP to Cas9-sgRNA complexes were prepared and measured for their sizes and charges (Table 1). Dynamic light scattering (DLS) measurements showed that the average hydrodynamic sizes of EP, Cas9 and Cas9-sgRNA (1:1) were 0.83 ± 0.05 nm, 7.8 ± 0.9 nm and 14.4 ± 1.0 nm, respectively. The hydrodynamic size of the CriPs consisting of Cas9-sgRNA-EP (1:1:20) was 376.8 ± 20.2 nm. The sizes of the CriPs were further increased when coated with more EPs, suggesting that each particle contains multiple Cas9-sgRNA complexes associated with EP peptides. The sizes of CriPs remained unchanged with dilution in the Dulbecco’s Modified Eagle’s Medium (DMEM) which mimics the *in vitro* cell culture conditions, suggesting the stability of the particles in the culture media.

**Table 1.** Size and charge measurements of particles.

|  | Size<br>(nm) | Charge (mV) |
| --- | --- | --- |
| **EP** | **$0.83 \pm 0.05$** | **$18.0 \pm 5.5$** |
| **Cas9** | **$7.8 \pm 0.9$** | **$13.9 \pm 6.8$** |
| **Cas9-sgRNA (1:1)** | **$14.4 \pm 1.0$** | **$-2.4 \pm 3.8$** |
| **Cas9-sgRNA-EP (1:1:20)** | **$376.8 \pm 20.2$** | **$6.0 \pm 4.4$** |
| **Cas9-sgRNA-EP (1:1:150)** | **$1240.9 \pm 243.8$** | **$14.6 \pm 6.3$** |
| **Cas9-sgRNA-EP (1:1:250)** | **$1238.7 \pm 34.8$** | **$20.7 \pm 6.0$** |
| **Cas9-sgRNA-EP (1:1:20) in DMEM media** | **$371 \pm 30.5$** | **N/A** |
| **Cas9-sgRNA-EP (1:1:150) in DMEM media** | **$1155.6 \pm 85.3$** | **N/A** |
| **Cas9-sgRNA-EP (1:1:250) in DMEM media** | **$1543.3 \pm 56.3$** | **N/A** |

Zeta potentials of the particles were also measured. Positive charges of +6.0 ± 4.4 mV, +14.6 ± 6.3 mV and +20.7 ± 6.0 mV were observed for the CriPs loaded with Cas9-sgRNA-EP with a molar ratio at 1:1:20, 1:1:150 and 1:1:250, respectively. The overall positive charge on the surface of the particles could facilitate cellular uptake by interacting with the negatively charged cell membranes.

### Deletion of *Gfp* in a macrophage cell line by CriPs

To investigate whether CriPs can efficiently delete genes *in vitro*, we first treated a macrophage cell line J774A.1 which stably expresses GFP (GFP-J774A.1) with CriPs loaded with sgRNA targeting the *Gfp* gene (*Gfp* sgRNA). The targeting sgRNA directed to *Gfp* is designed to introduce an indel mutation that results in a frame shift and termination of *Gfp* translation. CriPs were incubated with GFP-J774A.1 cells and the loss of GFP signal was determined by flow cytometry. 7-Amino-Actinomycin D (7-AAD) staining was used to distinguish among viable cells and dead cells. Percentages of GFP-negative cells and GFP-positive cells were calculated from the live cells. At 48 hours post treatment, CriPs loaded with *Gfp* sgRNA (Cas9- *Gfp* sgRNA -EP) induced a shift of the GFP signal in 54.0% of the GFP-J774A.1 cells (Fig. 2A and Fig. S1). The shift of the GFP signal rather than the complete loss of GFP after 48 hours of treatment was due to the slow degradation of the GFP protein. On day 5 post treatment, CriPs loaded with *Gfp* sgRNA (Cas9- *Gfp* sgRNA -EP) caused a loss of the GFP signal in 55.2% of the GFP-J774A.1 cells, while cells treated with EP only, Cas9- *Gfp* sgRNA only or without any treatment remained GFP positive in virtually all the cells (Fig. 2B).

**Figure 2.**
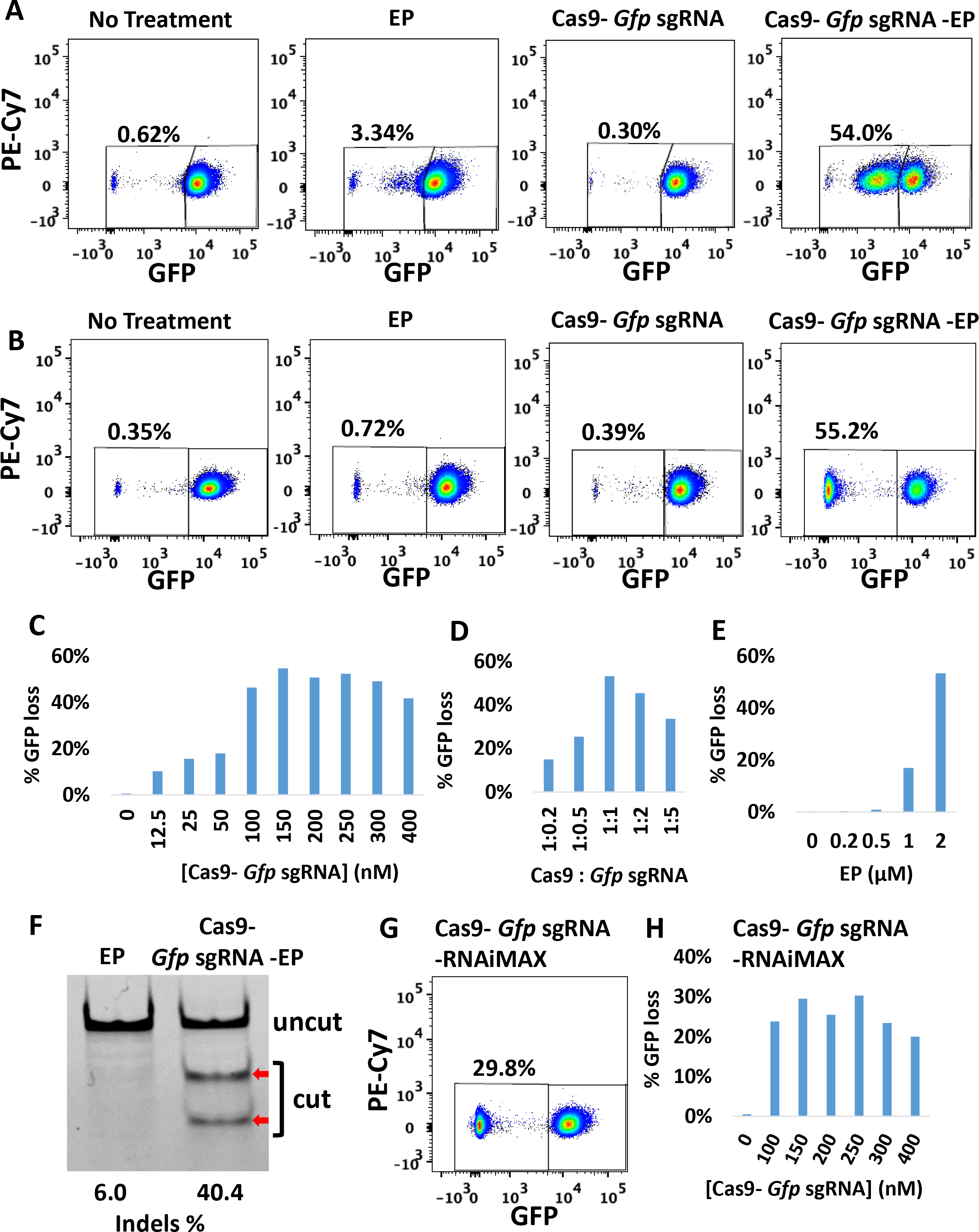
Efficient gene deletion achieved in GFP-J774A.1 cells following treatment with CriPs targeting Gfp. At 48 hours or on 5 days post treatment, flow cytometry and T7E1 assays were performed to measure the loss of GFP. (A) 48 hours post treatment, GFP loss measurements by flow cytometry. (B) 5 days post treatment, GFP loss measurements by flow cytometry. Cas9: 150 nM, *Gfp* sgRNA: 150 nM, EP: 2 μM. (C) Different concentrations of Cas9- *Gfp* sgRNA (1:1) with 2 μM of EP. (D) Different ratios of Cas9 (200 nM): *Gfp* sgRNA with 2 μM of EP. (E) Different concentrations of EP with 200 nM of Cas9- *Gfp* sgRNA. (F) Percent indels measurements in *Gfp* genomic DNA isolated from CriPs-treated cells vs EP treated cells by T7E1 assay. Uncut: 292 bp, Cut: 179 bp + 113 bp. Cas9: 200 nM, *Gfp* sgRNA: 200 nM, EP: 2 μM. (G) Flow cytometry data of GFP expressing J774A.1 cells treated with RNAiMAX mediated delivery of Cas9- *Gfp* sgRNA. Cas9: 150 nM, *Gfp* sgRNA: 150 nM. (H) Different concentrations of Cas9- *Gfp* sgRNA (1:1) with RNAiMAX (3 μl).

To optimize the gene deletion efficiency of CriPs in GFP-J774A.1 cells, we first determined the relationship between CriP dose and response by changing the concentrations of the Cas9- *Gfp* sgRNA complexes. GFP loss was observed in about 50% of the GFP-J774A.1 cells treated within the range of 100 nM to 300 nM of the Cas9- *Gfp* sgRNA complexes (Fig. 2C). We next tested the effect of the Cas9-sgRNA molar ratio on the gene deletion efficiency. A maximum loss of GFP in 53.3% of the GFP-J774A.1 cells was observed in response to CriPs loaded with a 1:1 molar ratio of Cas9 and *Gfp* sgRNA (Fig. 2D). By assessing the effect of EP concentration on *Gfp* deletion efficiency, a maximum GFP loss in 53.4% of the GFP-J774A.1 cells was observed with 2 μM of EP (Fig. 2E), which is the highest concentration of EP that was not associated with detectable cytotoxicity (Fig. S3).

To confirm the loss of GFP in the GFP-J774A.1 cells was due to genome engineering by the CriPs targeting the *Gfp* gene, the frequency of indel mutations in the *Gfp* genomic locus was determined by T7E1 assay. The cells treated with CriPs targeting *Gfp* (Cas9- *Gfp* sgRNA -EP) showed 40.4% of indels in the *Gfp* genomic locus, while the cells treated with EP alone showed only background noise (Fig. 2F). T7E1 assay usually underestimates the mutation frequencies due to the efficiency of the polymerase chain reaction (PCR) and the cleavage property of the T7 Endonuclease I(53). T7E1 recognizes and cleaves non-perfectly matched DNA at the first, second or third phosphodiester bond that is upstream of the mismatch. T7E1 cleaves heteroduplex DNA but does not recognize homozygous mutations, completely ignores single polymorphisms (SNPs) and also tends to miss small indels(53).

### Comparison of CriPs versus Lipofectamine^®^ RNAiMAX *in vitro*

To compare the gene deletion efficiency by CriPs versus Cas9-sgRNA delivered by commercially available transfect agents, we treated the GFP-J774A.1 cells with either CriPs loaded with *Gfp* sgRNA or Cas9- *Gfp* sgRNA plus Lipofectamine^®^ RNAiMAX. At two time points, 48 hours or 5 days post treatment of Cas9- *Gfp* sgRNA -RNAiMAX, a GFP shift or loss was observed as determined by flow cytometry in 29.4% and 29.8% of GFP-J774A.1 cells, respectively (Fig. 2G and Fig. S2). Cas9- *Gfp* sgRNA delivered by Lipofectamine^®^ RNAiMAX showed less *Gfp* gene deletion efficiency compared to CriPs which showed a GFP loss in over 50% of cells (Fig. 1B). To optimize the gene deletion efficiency of Cas9- *Gfp* sgRNA delivered by Lipofectamine^®^ RNAiMAX in GFP-J774A.1 cells, the concentrations of the Cas9- *Gfp* sgRNA complexes (Fig. S2B & S2D) and the molar ratio of the Cas9- *Gfp* sgRNA complexes were optimized (Fig. S2E & S2F). A maximum loss of GFP was observed in 20-30% of the GFP-J774A.1 cells treated with the range of 100 nM to 300 nM of the Cas9- *Gfp* sgRNA complexes at molar ratios of 1:1 to 1:5. These experiments suggest that *Gfp* gene deletion by Cas9- *Gfp* sgRNA delivered using Lipofectamine^®^ RNAiMAX was not as efficient as the CriPs in GFP-J774A.1 cells (Fig. 1A-E and Fig. S1).

### *Gfp* deletion in primary white pre-adipocytes from GFP mice

To investigate whether CriPs can efficiently delete genes in primary pre-adipocytes *in vitro*, primary pre-adipocytes were isolated from GFP transgenic heterozygous mice and treated with CriPs loaded *Gfp* sgRNA or control CriPs loaded with a control, non-targeting sgRNA (Control sgRNA). CriPs were formulated in two different buffer systems, NEBuffer 3 and phosphate-buffered saline (PBS). CriPs were then incubated with primary GFP white pre-adipocytes. On day 5 post treatment, the loss of GFP signal was determined by flow cytometry. Formulated in NEBuffer 3, CriPs loaded with *Gfp* sgRNA (Cas9- *Gfp* sgRNA-EP) induced a loss of the GFP signal in 48.5% of the GFP pre-adipocytes, while the control CriPs (Cas9- Control sgRNA -EP) showed only a background noise of 0.84% of the negative GFP pre-adipocytes (Fig. 3A). To optimize the EP dose that causes the maximum gene deletion efficiency of *Gfp* by CriPs formulated in NEBuffer 3 in the primary GFP pre-adipocytes, the relationship between EP dose and response was determined. A maximum GFP loss was achieved in about 43.7% ± 3.0% of the GFP pre-adipocytes when the Cas9- *Gfp* sgRNA complexes (100 nM) were coated with 30 μM of the EP (Fig. 3B). Interestingly, the loss of GFP was not complete after five days post transfection, as flow cytometry analysis showed a mixed population of cells that either displayed a complete loss of GFP, a shift in GFP loss, or no GFP loss. These variations can possibly be explained by the fact that the GFP transgenic mice likely have multiple *Gfp* transgenes inserted into their genome(54).

**Figure 3.**
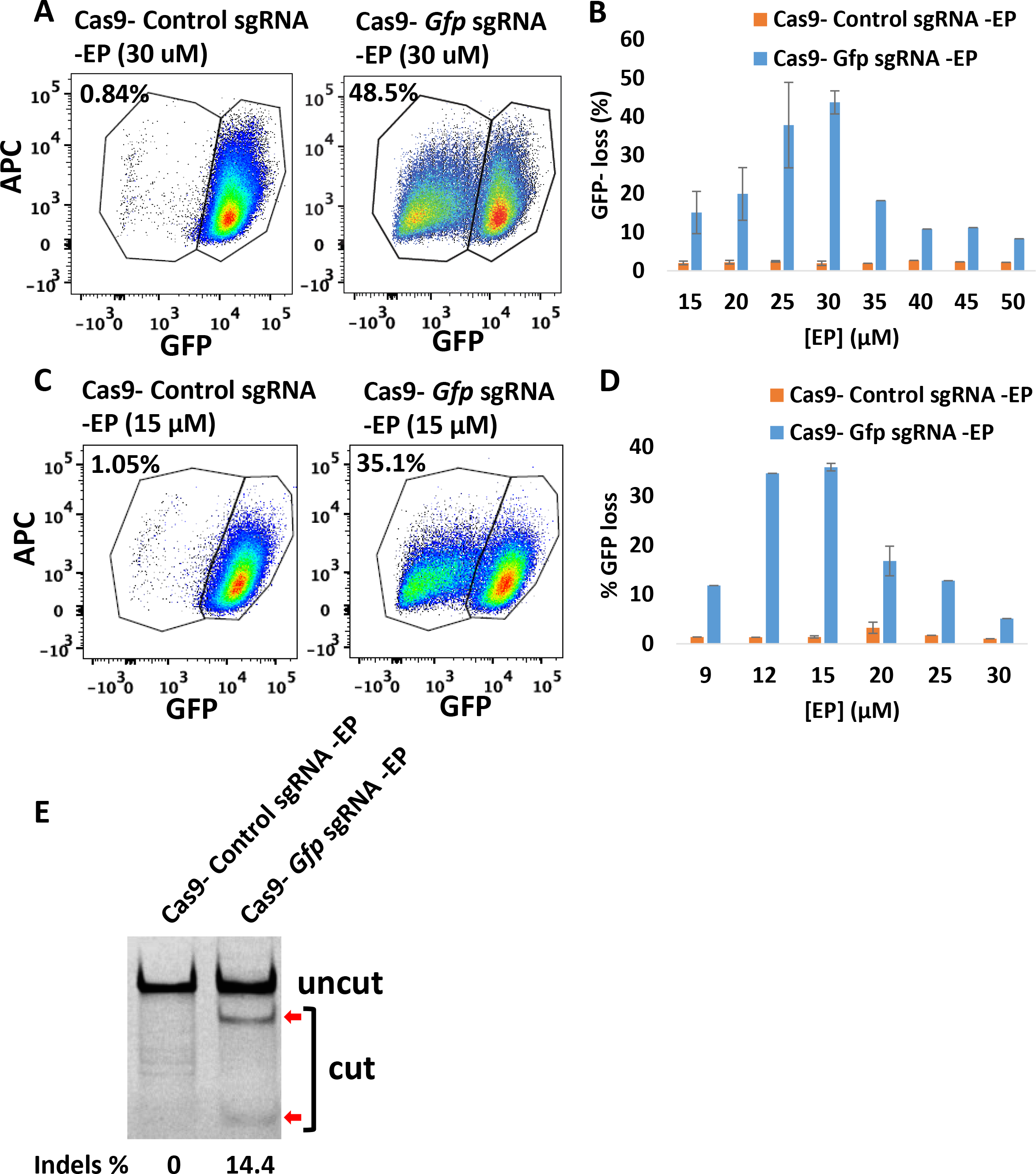
Efficient gene deletion achieved following the treatment with CriPs targeting *Gfp* in primary GFP pre-adipocytes isolated from GFP mice. (A, C) Flow cytometry data of primary GFP pre-adipocytes treated with CriPs formulated in (A) NEBuffer 3 or (C) PBS. Cas9-sgRNA: 100 nM. (B, D) Different concentrations of EP with 100 nM of Cas9-sgRNA formulated in (B) NEBuffer 3 or (D) PBS. (E) Percent Indels measurements by T7E1 assay in genomic DNA isolated from primary GFP pre-adipocytes. Uncut: 401 bp, Cut: 288 bp + 113 bp.

When CriPs were formulated in PBS, a loss of the GFP signal in 35.1% of the GFP pre-adipocytes was observed when treated with CriPs targeting *Gfp* (Cas9- *Gfp* sgRNA -EP), while the control CriPs (Cas9- Control sgRNA -EP) showed a background of 1.05% loss (Fig. 3C). The dose-response relationship of EP indicated a maximum GFP loss in about 35.8% ± 0.8% of the primary GFP pre-adipocytes treated with the CriPs loaded with the Cas9- *Gfp* sgRNA complexes (100 nM) coated with 15 μM of the EP (Fig. 3D). Thus, the gene deletion efficiency is dependent on the formulations of the CriPs in different buffer systems coated with different doses of EP.

To confirm the loss of the GFP in the primary GFP pre-adipocytes was due to genome engineering by the CriPs targeting the *Gfp* gene, we performed T7E1 assay to determine the mutation frequency of indels in the *Gfp* genomic locus. Compared to the cells treated with control CriPs, cleaved products were observed in the cells treated with CriPs targeting *Gfp* (Cas9- *Gfp* sgRNA -EP) with a mutation frequency of 14.4% (Fig. 3E). This is likely due to the multiple copies of GFP transgenes inserted into the genome(54) as well as an underestimation caused by T7E1 assay(53). These results indicate that efficient genome targeting can be achieved by treatment with CriPs in primary pre-adipocytes *in vitro*.

### Deletion of *Gfp* in PECs isolated from GFP transgenic mice

To determine the gene deletion efficiency using CriPs in primary GFP PECs, primary PECs that stably express GFP were isolated from GFP transgenic heterozygous mice which had been injected with thioglycollate broth. The PECs were treated with CriPs loaded with *Gfp* sgRNA or control CriPs loaded with Control sgRNA. On day 5 post treatment, the loss of GFP signal was determined by flow cytometry. Compared to control CriPs that showed a background shift of 2.88% (Fig. 4A), CriPs loaded with *Gfp* sgRNA (Cas9- *Gfp* sgRNA -EP) showed a decrease of the GFP signal in 40.7% of the GFP-expressing PECs (Fig. 4B). The relationship between Cas9-sgRNA dose and response indicated a maximum GFP loss of about 35.2% ± 4.0% in the GFP-expressing PECs when the CriPs were loaded with 100 nM of the Cas9- *Gfp* sgRNA complexes coated with 2 μM of the EP (Fig. 4C). T7E1 assay showed a mutation frequency of 32.8% in the *Gfp* genomic DNA isolated from the cells treated with CriPs targeting *Gfp*, compared to a background of 2.4% in the cells treated with the control CriPs (Fig. 4D).

**Figure 4.**
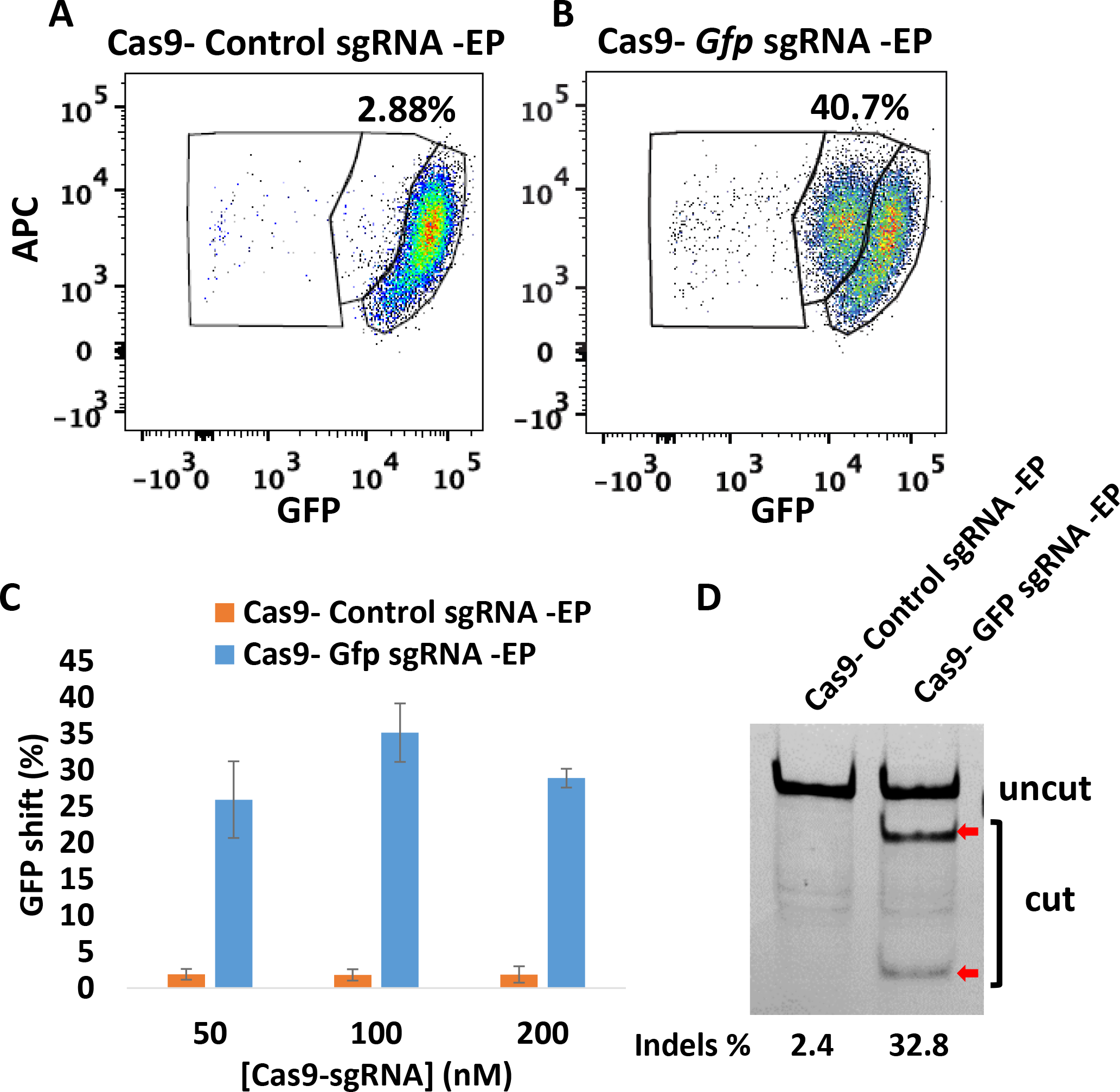
Efficient gene deletion achieved following treatment with CriPs targeting *Gfp* in primary PECs isolated from thioglycollate-injected GFP mice. 5 days post treatment, flow cytometry and T7E1 assay were performed to measure the loss of GFP. Flow cytometry data of primary GFP PECs treated with (A) CriPs with a control, non-targeting sgRNA sequence (Cas9-Control sgRNA -EP) and (B) CriPs targeting GFP (Cas9- *Gfp* sgRNA -EP). Cas9: 100 nM, sgRNA: 100 nM, EP: 2 μM. (C) Different concentrations of Cas9-sgRNA (1:1) with 2 μM of EP. (D) Percent Indels measurements in *Gfp* genomic DNA isolated from CriPs-treated primary GFP PECs by T7E1 assay. Uncut: 401 bp, Cut: 288 bp + 113 bp. Cas9: 100 nM, sgRNA: 100 nM, EP: 2 μM.

### Deletion of *Gfp* by delivery of CriPs *in vivo* to GFP mice

To determine the ability of CriPs to systemically deliver Cas9-sgRNA RNPs *in vivo* to achieve gene deletion, we administrated CriPs to mice by i.p. injections. Shown in Figure 5A, GFP transgenic heterozygous mice were i.p. injected daily for 5 days with CriPs targeting *Gfp* (CriPs-*Gfp* sgRNA) or with control CriPs (CriPs- Control sgRNA). On day 6, mice were sacrificed and PECs were collected and plated in cell culture. On day 13, flow cytometry was performed to measure the loss of GFP signal. Deep sequencing was also performed to detect the indels in the *Gfp* genomic locus. Measured by flow cytometry, a loss of the GFP signal was observed in the range of 2.9%-13.9% of the PECs (average 5.70% ± 1.07%) isolated from ten GFP transgenic heterozygous mice injected with CriPs- *Gfp* sgRNA. This was significantly greater than the less than 1% GFP loss we observed in the CriPs- Control sgRNA treated mice (average 0.74% ± 0.05%) (Fig. 5B and 5C).

**Figure 5.**
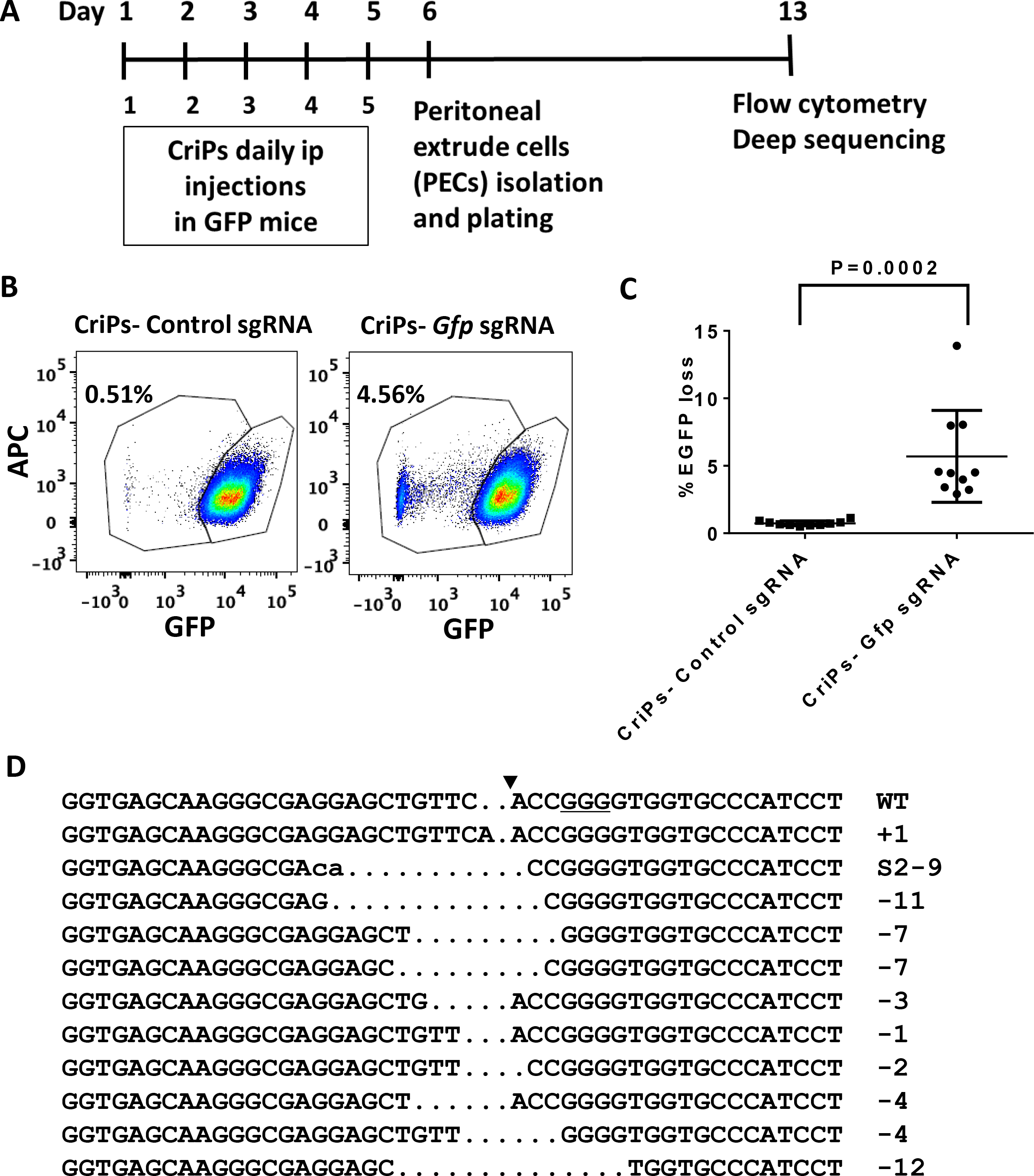
Efficient gene deletion achieved by i.p. injections of CriPs targeting *Gfp* in GFP mice. (A) Timeline of i.p. administration of CriPs to GFP mice. GFP transgenic mice were i.p. injected daily for 5 days with CriPs targeting *Gfp* (CriPs- *Gfp* sgRNA) or control CriPs (CriPs- Control sgRNA). On day 6, mice were sacrificed and PECs were collected and plated in cell culture. On day 13, flow cytometry and deep sequencing were performed to measure the loss of GFP. (B) Flow cytometry data showing GFP loss in mice injected with CriPs- *Gfp* sgRNA or CriPs-Control sgRNA. (C) Quantification of flow cytometry data. 3-8 week old GFP male C57BL/6 mice, n = 10 (D) Indels in the *Gfp* locus of CriPs- *Gfp* sgRNA injected samples by deep sequencing. DNA sequences of the *Gfp* wild-type (WT) and mutants. PAM is underlined. The cleavage site is indicated by an arrowhead. The column on the right indicates the number of inserted (+) or deleted (−) bases or SNPs (S).

In order to confirm the deletion of GFP was due to genome engineering of *Gfp* by CriPs- *Gfp* sgRNA as well as to study the mutation composition of the insertions and deletions in the *Gfp* genomic locus, we performed deep sequencing on the PCR amplicons amplified from the genomic DNA isolated from PECs. Approximately 3% of the target sequences were mutated in the CriPs- *Gfp* sgRNA injected samples, corroborating the data on the GFP loss measured by flow cytometry shown in Figure 5B, while only a background noise of 0.02% was observed in the CriPs- Control sgRNA injected samples. To study the composition of the mutations in the *Gfp* genomic locus of the CriPs- *Gfp* sgRNA injected samples, the DNA sequences of the GFP wild type (WT) and the mutants were aligned (Fig. 5D). The mutants consisted of sequences with insertions and deletions of the bases as well as SNPs in the target site.

Taken together, significant gene deletion of the *Gfp* gene was achieved by i.p. injections of the CriPs targeting *Gfp in vivo* in the GFP transgenic heterozygous mice, as determined by both flow cytometry and deep sequencing. To the best of our knowledge, this is the first study demonstrating a simple non-viral genome editing system that delivers Cas9 protein loaded with sgRNA systemically *in vivo* to achieve significant genome engineering. This is likely an underestimate of the efficiency since GFP expressing transgenic mice are predicted to have multiple GFP transgenes inserted into their genome(54). A higher genome engineering efficiency would be expected when editing an endogenous target gene of interest that is present in two copies.

### Analysis of immune response in mice injected with CriPs

To examine the possibility of an immune response and systematic inflammation induced by CriPs, we i.p. injected WT mice daily for 5 days with CriPs loaded with a non-targeting sgRNA or PBS as a control. The systemic cytokine profile was analyzed by measuring cytokine levels (IL-1β, IL-4, IL-6, IL-10, IFNγ, TNFα) in the plasma before injection, and then 24 hours and 2 weeks after the first injection. The plasma cytokine levels (IL-10 and TNFα) in mice treated with LPS for 1.5 hours were also measured to serve as a positive control for the assay (Fig. 6D).

**Figure 6.**
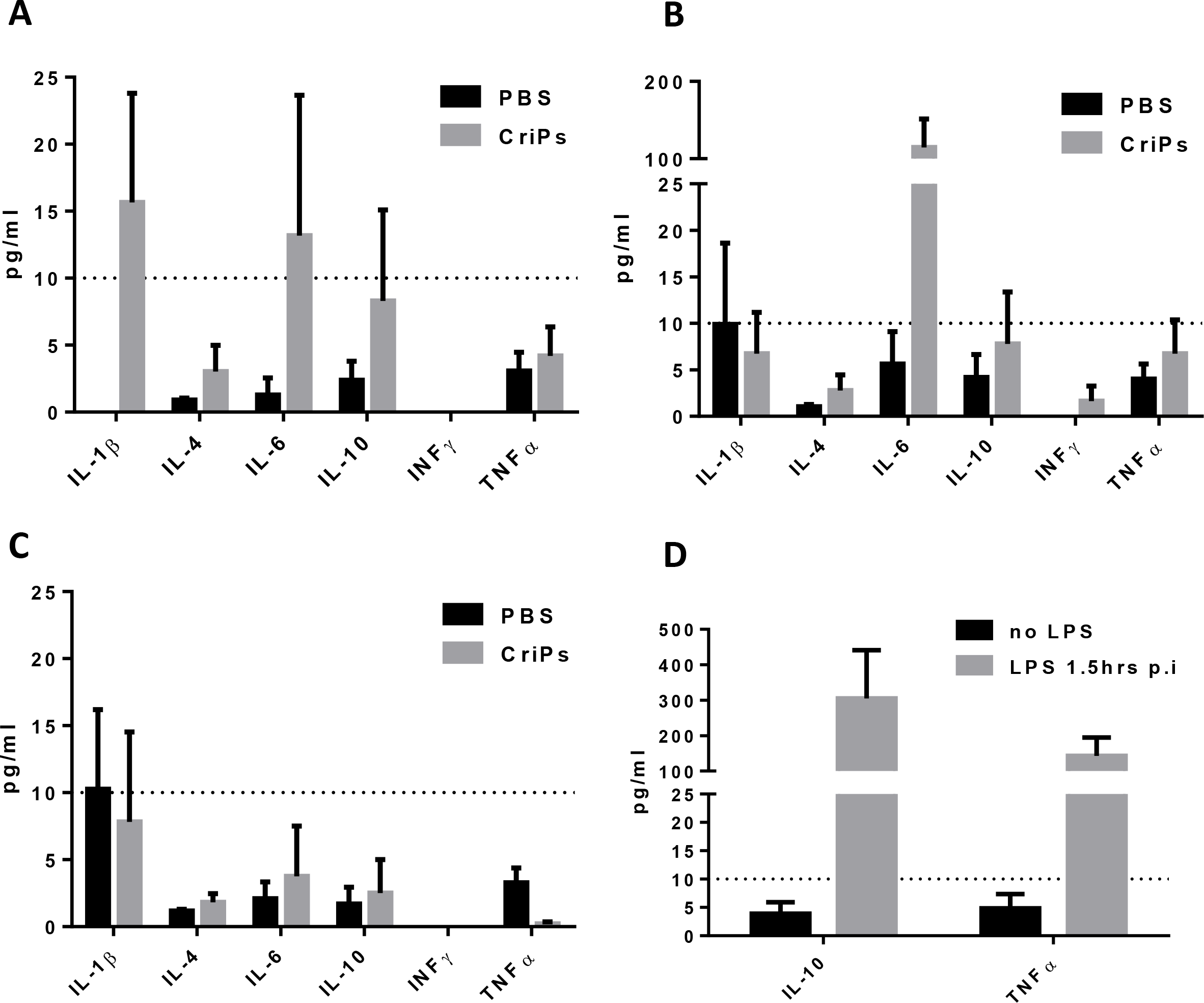
Plasma cytokine levels were measured in mice i.p. injected with CriPs *in vivo*. WT mice were i.p. injected daily for 5 days with PBS or CriPs loaded with a non-targeting sgRNA. Plasma was collected before injection, and then 24 hours and 2 weeks after the first injection. Plasma cytokine levels (IL-1β, IL-4, IL-6, IL-10, IFNγ, TNFα) were measured by Luminex. Plasma cytokine levels were measured (A) before injection (B) 24 hours after the first injection (C) 2 weeks after the first injection. (D) Plasma cytokine levels (IL-10 and TNFα) were also measured in the mice treated with LPS at 1.5 hours post injection (p.i.) or without LPS to serve as positive controls for the assay. Data are means ± sem (n = 3-4).

Shown in Figure 6A, low background levels of inflammatory cytokines were observed before i.p. injection of CriPs and PBS in mice. Systemic injections of CriPs did not cause an acute (24 hours) or a chronic (2 weeks) up-regulation of inflammatory cytokines such as IL-1β, IL-4, IL-10, IFNγ and TNFα in the plasma (Fig. 6B and 6C), suggesting the absence of a broad immune response upon CriPs administration under the conditions of our experiments. Interestingly, the level of IL-6 was increased in the plasma 24 hours after the first injection in mice treated with CriPs. IL-6 acts as both a pro-inflammatory cytokine and an anti-inflammatory myokine(55). It can be significantly elevated by many factors such as exercise(56). More importantly, IL-6 returned to a low background level when it was measured 2 weeks after the first injection. In addition, the low levels of all the plasma inflammatory cytokines measured 10 days after total 5 injections (2 weeks after the first injection) indicated that CriPs can be administrated multiple times systemically *in vivo* without chronic toxicity (Fig. 6C).

### Deletion of *Nrip1* in primary pre-adipocytes enhances “browning”

Targeting genes in adipocytes that suppress mitochondrial uncoupling and fatty acid oxidation is a potential strategy to enhance adipocyte “browning” and energy expenditure(37). Our laboratory has used RNAi-based screens in cultured adipocytes to identify such genes that control insulin sensitivity and energy metabolism(43). An exciting “hit” in these screens of several thousand genes was the nuclear co-repressor *Nrip1*(43), which was also discovered in independent studies in mice(45). NRIP1 interacts with nuclear receptors to suppress their activities to regulate genes that control glucose utilization, mitochondrial function, fatty acid oxidation and secretion of beneficial factors(43). We and others have found that the depletion of NRIP1 in adipocytes and in mice increased fatty acid oxidation, mitochondrial respiration, insulin sensitivity and glucose tolerances(43,45,57). Therefore, NRIP1 presents a potential therapeutic target for type 2 diabetes and other metabolic disorders.

In order to investigate whether the *Nrip1* gene can be efficiently deleted in adipocytes and whether the depletion of *Nrip1* can lead to the enhanced expression of genes associated with energy expenditure, such as UCP1, we isolated primary white pre-adipocytes from WT mice and treated them with CriPs loaded with each of four different sgRNAs targeting *Nrip1* (*Nrip1* sgRNA 1, *Nrip1* sgRNA 2, *Nrip1* sgRNA 3, *Nrip1* sgRNA 4) or control groups (CriPs-Control sgRNA, Cas9-EP, EP only and non-treated). The treated pre-adipocytes were then differentiated to mature adipocytes using a differentiation cocktail. On day 8 post differentiation, cells were collected to measure both the frequency of indels in the *Nrip1* genomic locus by T7E1 assay and the expression of UCP1 by RT-PCR.

Indicated by T7E1 assay (Fig. 7A), four CriP formulations, each loaded with a different sgRNA targeting *Nrip1*, showed indels in the *Nrip1* genomic locus with different degrees of mutation frequencies. CriPs loaded with *Nrip1* sequence #3 (*Nrip1* sgRNA 3) demonstrated the highest mutation frequency of 43.8% in the *Nrip1* genomic locus. More importantly, the deletion of the *Nrip1* gene in the white adipocytes caused the “browning” of cells, evidenced by increased expression of the uncoupling protein UCP1. Indeed, UCP1 was significantly increased in response to all four CriP formulations loaded with different sgRNAs targeting *Nrip1*. Adipocytes treated with CriPs loaded with the *Nrip1* sgRNA 3 that was most potent in deleting the *Nrip1* gene measured by T7E1 assay (Fig. 7A), elicited the most marked increase in UCP1 expression (Fig. 7B).

**Figure 7.**
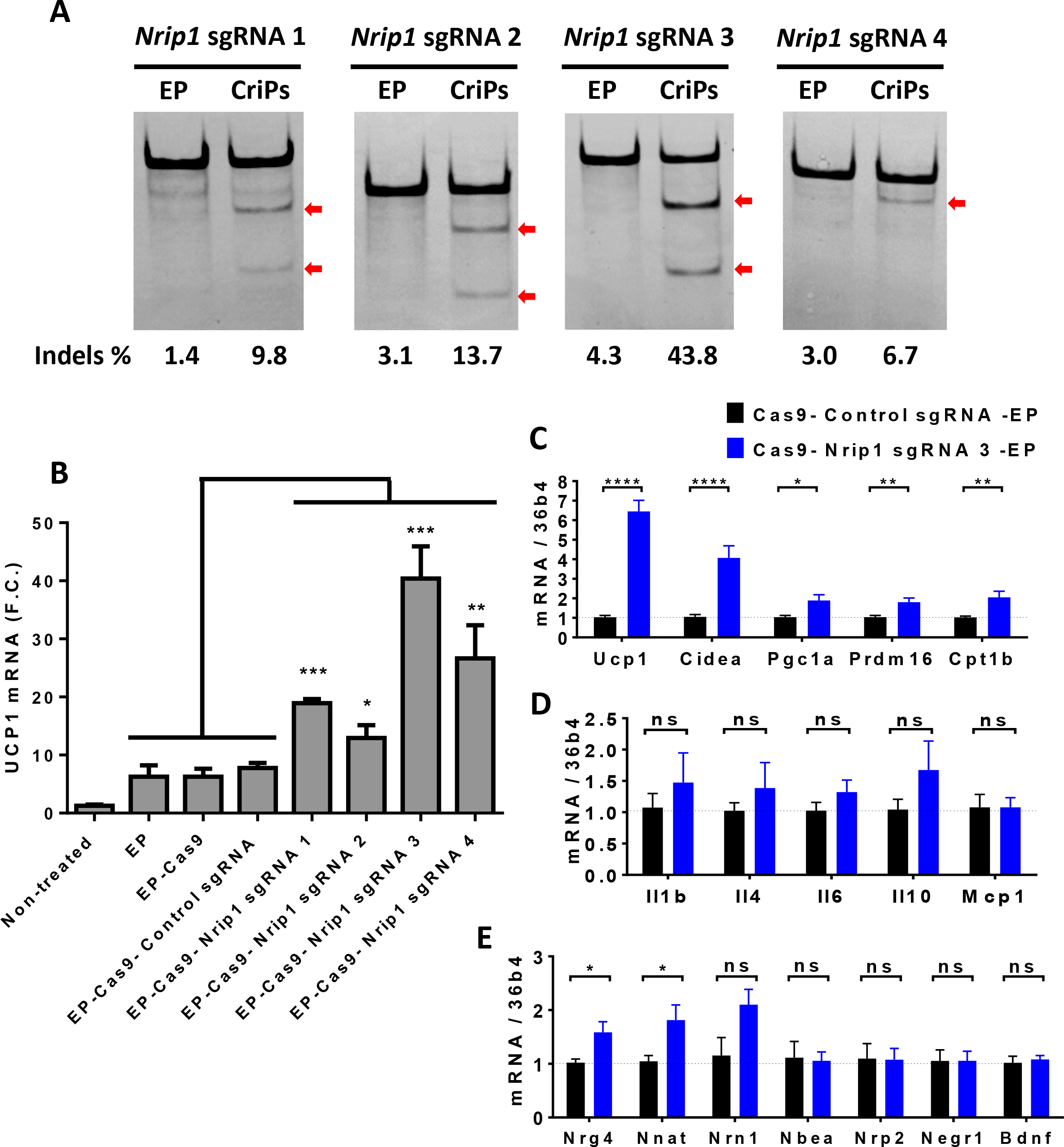
CriPs-mediated *Nrip1* deletion in white adipocytes increases the expression of UCP1. Primary pre-adipocytes were treated with CriPs with each of four different sgRNAs targeting *Nrip1* (*Nrip1* sgRNA 1, *Nrip1* sgRNA 2, *Nrip1* sgRNA 3, *Nrip1* sgRNA 4) and controls. Pre-adipocytes were then differentiated into mature white adipocytes. Cas9-sgRNA: 100 nM, EP: 25 μM. (A) Indels detection in *Nrip1* genomic DNA was determined by T7E1 assay. *Nrip1* sgRNA 1: Uncut: 429bp, Cut: 283bp + 146bp; *Nrip1* sgRNA 2: Uncut: 334bp, Cut: 223bp + 111bp; *Nrip1* sgRNA 3: Uncut: 420bp, Cut: 150bp + 270bp; *Nrip1* sgRNA 4: Uncut: 381bp, Cut: 307bp + 74bp. (B) UCP1 expression was measured by RT-PCR. (C, D, E) With the treatment of Cas9-*Nrip1* sgRNA 3 -EP and Cas9- Control sgRNA -EP, gene expression was measured by RT-PCR. (C) thermogenic genes (D) inflammatory genes (E) neurotropic factors

The expression of other thermogenic genes, inflammatory genes and neurotropic factors were also measured by RT-PCR in the white adipocytes that had been treated with Cas9- *Nrip1* sgRNA 3 -EP and Cas9- Control sgRNA -EP. Thermogenic genes involved in adipocyte “browning” (*Ucp1*, *Cidea*, *Pgc1α*, *Prdm16*, *Cpt1b*) were increased with the treatment of CriPs loaded with *Nrip1* sgRNA 3, compared to the control CriPs (Fig. 7C). Neurotropic factors (*Nrg4*, *Nnat*, *Nrn1*) were also increased with the treatment of CriPs- *Nrip1* sgRNA 3 (Fig. 7E), while inflammatory genes (*Il1β*, *Il4*, *Il6*, *Il10*, *Mcp1*) were not changed between CriPs- *Nrip1* sgRNA 3 and CriPs- Control sgRNA groups (Fig. 7D). Overall, these results indicate that efficient *Nrip1* gene deletion was achieved in white adipocytes treated with CriPs targeting *Nrip1*, and the deletion of the *Nrip1* gene by CriPs converts the white adipocytes to a more “brown” adipocyte phenotype known to be beneficial to whole-body metabolism in mice.

### Off-target effects of CriPs targeting *Nrip1* in adipocytes

Primary pre-adipocytes were treated with CriPs loaded with *Nrip1* sgRNA 3 that caused the highest mutation frequency in the *Nrip1* genomic locus and most increased UCP1 in differentiated adipocytes, as indicated in Figure 7. Differentiated adipocytes were collected to measure the mutation frequencies in the on-target *Nrip1* genomic locus and the off-target genomic sites. All the off-target sequences contained three mismatch bases compared to the on-target sequence and were located in the intergenic regions. Determined by T7E1 assay, no obvious off-target effects in the adipocytes treated with CriPs targeting *Nrip1* were observed compared to the non-treated cells (Fig. 8).

**Figure 8.**
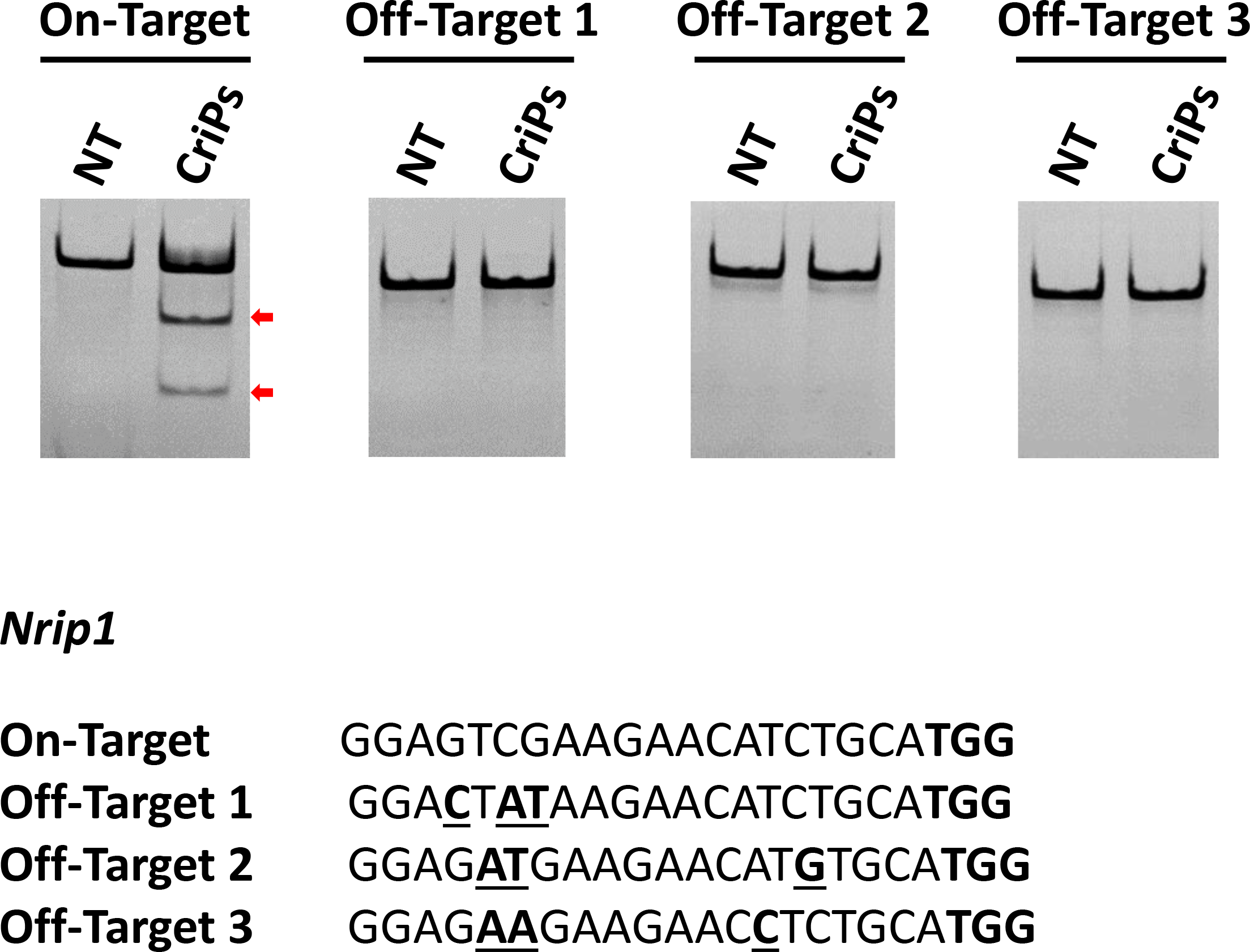
Determination of off-target effects of CriPs targeting *Nrip1* by T7E1 assay. Primary pre-adipocytes were treated with CriPs with *Nrip1* sgRNA 3. Pre-adipocytes were then differentiated into mature white adipocytes. Top off-target candidate sites were determined by the CHOPCHOP program. Off-target effects were determined by T7E1 assay. Expected DNA bands cleaved by T7E1: On-target: Uncut: 420bp, Cut: 270bp+150bp; Off-target 1: Uncut: 386bp, Cut: 283bp+103bp; Off-target 2: Uncut: 387bp, Cut: 229bp+158bp; Off-target 3: Uncut: 352bp, Cut: 182bp+170bp. TGG in bold is the PAM site. Mismatch sites are underlined and in bold.

## DISCUSSION

A key challenge to realizing the potential of CRISPR-Cas9-based therapeutics is the assurance of safe and effective delivery to target cells(6, 9). The major advance of this study is the development of a novel and simple system, CriPs, to efficiently deliver CRISPR-Cas9-based reagents *in vitro* and *in vivo*. CriPs consist of three components ‑ Cas9 protein, sgRNA targeting genes of interest and the EP peptide, which facilitates transport of large cargoes across cell membranes (Fig 1). As a therapeutic approach, the direct delivery of Cas9 in protein form enables the swiftest gene editing as there is no need for transcription or translation of the nuclease(7). Direct delivery of Cas9 protein offers advantages over plasmid delivery and viral delivery, which can display uncontrolled integration of DNA segments into the host genome, unwanted immune responses to the plasmids and virus, and limited packaging capacity(6). It has been shown that the Cas9 protein introduced into cells rapidly degrades within 24 hours(52), eliminating immune responses using an *ex vivo* therapeutic approach where implantation of engineered cells is performed days after gene deletion. Since CriPs-mediated gene editing lasts for the lifespan of a cell, for example up to 10 years for adipocytes(58), infrequent administration of engineered cell implants would be required to maintain therapeutic efficacy.

Characterized by DLS (Table 1), the CriPs composed of Cas9-sgRNA-EP (1:1:20) showed a hydrodynamic size of 376.8 ± 20.2 nm with a positive zeta potential. Compared with Cas9-sgRNA (1:1) alone with a hydrodynamic size of 14.4 ± 1.0 nm, the increase of size indicates the formation of CriPs containing multiple Cas9-sgRNA complexes associated with EP peptides. The overall positive charge on the surface of the particles is predicted to facilitate the cellular uptake by interacting with the negatively charged cell membranes. The sizes and the positive charges of CriPs were further increased when Cas9-sgRNA nanocomplexes were exposed to higher EP concentrations. Interestingly, the unchanged size of CriPs when exposed to DMEM media suggests stability of the particles in culture media.

The present study demonstrates that CriPs facilitate efficient gene deletion of the proof-of-concept gene *Gfp* in multiple cell types including GFP-J774A.1 cells (Fig. 2), GFP-PECs (Fig. 4) and primary GFP-expressing pre-adipocytes (Fig. 3). We observed GFP loss in about 50% of cells treated with CriPs targeting *Gfp*, detected by flow cytometry analysis (Fig. 2A-E, 3A-D, 4A-C) and confirmed by identification of indels in the *Gfp* genomic locus by T7E1 assay (Fig. 2F, 3E, 4D). We found that the EP concentration that is optimal for maximal gene deletion is highly cell type dependent (Fig. 2E, 3B, 3D). Primary pre-adipocytes require a much higher dose of EP to achieve maximum gene deletion compared to macrophages. On the contrary, high doses of EP are toxic to macrophages but not to primary pre-adipocytes (Fig. S3). Gene deletion efficiency also depends on the formulations of the CriPs in different buffer systems (Fig. 3A-D), possibly due to salts that affect the activity of the Cas9 protein and the electrostatic association of the Cas9-sgRNA nanocomplex and EP. In addition, CriPs demonstrate higher gene deletion efficiency in GFP-J774A.1 cells compared to the Cas9-sgRNA delivered by commercially available Lipofectamine^®^ RNAiMAX (Fig. 2G and 2H).

Although many delivery systems have been used to edit genes by Cas9-sgRNA RNPs in cell culture(30, 33), primary cells(15, 19) and local areas of animals such as inner ear (4, 23), skin(18), tumor(28), muscle(29) and brain(34), systemic delivery of Cas9-sgRNA complexes using a fully non-viral system has not been previously demonstrated. To determine the ability of CriPs to delete genes *in vivo* by systemic delivery, we performed 5 daily i.p. injections of CriPs targeting *Gfp* (CriPs- *Gfp* sgRNA) or control CriPs (CriPs- Control sgRNA) in GFP transgenic heterozygous mice (Fig. 5A). These mice showed a loss of the GFP signal in 2.91%-13.90% of the PECs when injected with CriPs- *Gfp* sgRNA, which was significantly higher than the CriPs-Control sgRNA treated group (average 0.74% ± 0.05%) (Fig. 5B and 5C). We also confirmed that about 3% of the target sequences were mutated in the CriPs- *Gfp* sgRNA injected animals, and the mutants were composed of insertions and deletions of the bases as well as SNPs (Fig. 5D). This is likely an underestimate of the targeting efficiency since GFP transgenic mice likely have multiple *Gfp* transgenes inserted into their genome(54). Taken together, our work demonstrates a simple non-viral genome editing system that delivers Cas9-sgRNA RNPs systemically *in vivo* to achieve significant gene deletion.

Gene editing or deletion in adipose tissue to enhance adipose tissue energy expenditure and fatty acid oxidation through a “browning” process presents a potential therapeutic approach to alleviate obesity and type 2 diabetes(37, 40). Brown adipocytes not only generate heat but also secrete beneficial factors that enhance glucose tolerance in mice(38, 42). We demonstrate the utility of CriPs-mediated gene deletion of *Nrip1* in white adipocytes, demonstrating an induced “browning” phenotype with remarkably enhanced expression of UCP1, known to uncouple mitochondrial respiration, activate fatty acid oxidation and improve glucose tolerance (Fig. 7). Using CriPs targeting *Nrip1* in the white adipocytes, we observed a highest mutation frequency of 43.8% in the *Nrip1* genomic locus with *Nrip1* sgRNA 3 measured by T7E1 assay (Fig. 7A). Adipocytes treated with the CriPs- *Nrip1* sgRNA 3 that were most potent in deleting the *Nrip1* gene, demonstrated the most marked increase in UCP1 expression (Fig. 7B). Other thermogenic genes (*Ucp1*, *Cidea*, *Pgc1α*, *Prdm16*, *Cpt1b*) and neurotropic factors (*Nrg4*, *Nnat*, *Nrn1*) were also increased (Fig. 7C and 7E). In addition, off-target effects in adipocytes treated with CriPs-*Nrip1* sgRNA 3 were not detected by T7E1 assay (Fig. 8). Thus the CriPs loaded with sgRNA targeting *Nrip1* provide significant potential for therapeutic development to alleviate metabolic disease.

## EXPERIMENTAL PROCEDURES

### Materials

All chemicals were purchased from Sigma-Aldrich (St. Louis, MO, USA) unless otherwise specified and were used as received. Cas9 protein was purchased from PNA BIO, INC. (Newbury Park, CA, USA). DNA oligonucleotides were purchased from Integrated DNA Technologies Inc. (Coralville, IA, USA). MEGAshortscript T7 Transcription Kit, Lipofectamine^®^ RNAiMAX, Vybrant MTT cell proliferation assay and Platinum™ Taq DNA Polymerase High Fidelity kit were purchased from ThermoFisher Scientific (Waltham, MA, USA). BsaI, DraI, T7E1 and NEBuffer 3 were obtained from New England Biolabs Inc. (Ipswich, MA, USA). pUC57-sgRNA expression vector was purchased from Addgene (plasmid # 51132) (Cambridge, MA, USA). Endo-Porter (EP) was purchased from Gene Tools (Philomath, OR, USA). QIAquick PCR Purification Kit was purchased from Qiagen Inc. (Valencia, CA, USA). 4-20% Mini-Protean TBE Gel was purchased from Bio-Rad Laboratories (Hercules, CA, USA). 7-AAD was purchased from BD Biosciences (San Jose, CA, USA). Insulin was purchased from Cell Application (San Diego, CA, USA) Dexamethasone, 3-isobutyl-1-methylxanthine (IBMX), indomethacin and lipopolysaccharide (LPS) were purchased from Sigma (St. Louis, MO, USA). Rosiglitazone was purchased from Cayman Chemical (Ann Arbor, MI, USA).

### Preparation of sgRNA template and synthesis of sgRNA

sgRNA sequences were designed using sgRNA Designer(59) developed by Broad Institute and CHOPCHOP program(60, 61) developed by Harvard University. Templates for sgRNAs were generated by inserting annealed complementary oligonucleotides with the sgRNA sequences to the pUC57-sgRNA expression vector encoding a T7 promoter. The sgRNA templates were linearized by Dra I and transcribed *in vitro* using the MEGAshortscript T7 Transcription Kit according to manufacturer’s instruction. Transcribed sgRNA was resolved on a 10% denaturing urea-Polyacrylamide gel electrophoresis to check the size and purity.

### Preparation of the CriPs

Purified bacterial Cas9 protein is processed to remove endotoxin and other contaminants and is used for loading of sgRNA. The powder of Cas9 protein was resuspended in water with 20% glycerol. Cas9 protein and sgRNA were mixed in NEBuffer 3 (100 mM NaCl, 50 mM Tris-HCl, 10 mM MgCl2, 1 mM DTT, pH 7.9) purchased from New England Biolabs Inc. or PBS at 37 °C for 10 minutes to form nano-size complexes. The loaded Cas9-sgRNA nanocomplexes are then complexed with EP in PBS at room temperature for 15 minutes to form the final CriPs.

### Characterization of the CriPs

The size of CriPs was determined by DLS (laser wavelength 633 nm) using a Malvern Zetasizer Nano-ZS particle size analyzer (Malvern Instruments, Worcestershire, UK). Solvents and buffers were filtered through 0.22 μm filters before sample preparation. EP (1mM), Cas9 (1 μM), Cas9-sgRNA (1 μM) and CriPs (1 μM) with different ratios of EP were measured for sizes upon the absence or presence of DMEM (GE Healthcare Life Sciences, Pittsburgh, PA). Zeta potentials of CriPs were also determined with a Malvern Zetasizer Nano-ZS using a Universal ‘Dip’ Cell Kit. Solvents and buffers were filtered through 0.22 μm filters before sample preparation. A suspension of samples was diluted in 20 mM of HEPES (4-(2-hydroxyethyl)-1-piperazineethanesulfonic acid) buffer for the measurement. Data were analyzed with the Dispersion Technology software (Malvern).

### Cell lines and culture

J774A.1 cells were acquired from ATCC (American Type Culture Collection, Manassas, VA). J774A.1 cells stably expressing GFP (a gift from Dr. H. Yang, University of Massachusetts Medical School, Worcester, MA) were maintained in DMEM supplemented with 10% (v/v) fetal bovine serum (FBS) (Atlanta Biologicals, Flowery Branch, GA), 100 μg/mL streptomycin and 100 units/mL penicillin (Thermo Fisher Scientific, Waltham, MA). Cell incubations were performed in a water-jacketed 37°C/5% CO_2_ incubator.

### *In vitro* CriPs treatment in GFP-J774A.1 cells

GFP-J774A.1 cells were plated in 12 well plates with 1×10^5^ cells per well overnight. Cells were treated with CriPs loaded with *Gfp* sgRNA (Cas9- *Gfp* sgRNA–EP) or controls such as Cas9-*Gfp* sgRNA without EP, EP only and non-treated. After 24 hours, media containing CriPs and controls was replaced with fresh culture media. At 48 hours or on day 5 post treatment, flow cytometry analysis was performed to measure the loss of GFP. 7-AAD staining was used to determine live cells and dead cells. The percentages of GFP-negative cells and GFP-positive cells were calculated from the live cells in the flow cytometry analysis. Indels in the *Gfp* genomic locus were measured by T7E1 assay.

### Transfection of GFP-J774A.1 with Cas9-sgRNA using RNAiMAX

GFP-J774A.1 cells were plated in 12 well plates with 1×10^5^ cells per well overnight. Cells were treated with Cas9- *Gfp* sgRNA complexes delivered by Lipofectamine^®^ RNAiMAX or controls such as Cas9- *Gfp* sgRNA without RNAiMAX, RNAiMAX only and non-treated. After 24 hours, cells were fed with fresh culture media. At 48 hours or on day 5 post treatment, flow cytometry analysis was performed to measure the loss of GFP. 7-AAD staining was used to determine live cells and dead cells. The percentages of GFP-negative cells and of GFP-positive cells were calculated from the live cells analyzed by flow cytometry.

### Animals

All mice were purchased from Jackson Laboratory. WT mice (C57BL/6J) and GFP transgenic heterozygous mice (C57BL/6-Tg(UBC-GFP)30Scha/J) were used for the experiments. Mice were housed on a 12-h light/dark schedule and had free access to water and food. All procedures involving animals were approved by the Institutional Animal Care and Use Committee at the University of Massachusetts Medical School.

### Isolation of primary PECs from GFP transgenic mice

Ten-week-old GFP transgenic heterozygous mice were i.p. injected with 4% thioglycollate broth (Sigma-Aldrich, St. Louis, MO). Five days following injection, the mice were sacrificed and the peritoneal cavity was washed with 5 mL of ice-cold PBS to isolate PECs. Peritoneal fluid was filtered through a 70 μm pore nylon mesh and centrifuged at 1200 rpm for 10 minutes. The pellet was first treated with red blood cell (RBC) lysis buffer (8.3 g of NH4Cl, 1.0 g of KHCO3 and 0.037 g of EDTA dissolved in 1 L of water) and resuspended in DMEM supplemented with 10% (v/v) FBS, 100 μg/mL streptomycin and 100 units/mL penicillin.

### *In vitro* CriPs treatment of primary GFP PECs

Primary GFP PECs isolated from GFP transgenic mice were plated in 12 well plates with 5×10^5^ cells per well overnight. Cells were treated with CriPs loaded with *Gfp* sgRNA or Control sgRNA. After 24 hours, cells were fed with fresh culture media. On day 5 post treatment, flow cytometry analysis was performed to measure the loss of GFP. 7-AAD staining was used to determine live cells and dead cells. The percentages of GFP negative cells and GFP positive cells were calculated from the live cells analyzed by flow cytometry. Indels in the *Gfp* genomic locus were measured by T7E1 assay.

### Culture of primary white pre-adipocytes from mice

Three-week-old GFP transgenic heterozygous mice or WT mice were sacrificed and inguinal subcutaneous fat pads were dissected out and placed into Hanks’ Balanced Salt solution (HBSS without Ca2+) with 3% bovine serum albumin (BSA). Tissues were minced with scissors to approximately 3-5 mm pieces. Tissues were digested in Collagenase D solution (2 mg/mL Collagenase D in HBSS with 3% BSA) in a 37 °C water bath shaker for 1 hour with short vortex every 10-15 minutes. Samples were inactivated with 10% FBS, filtered through a 100 μm mesh and centrifuged at 600g for 5 minutes. The stromal vascular fraction (SVF) pellet was resuspended in RBC lysis buffer for 5 minutes and centrifuged again at 600g for 5 minutes. Cell pellet was resuspened in DMEM/F12 media with 10% FBS and 1% streptomycin/penicillin, filtered through 40 μm mesh and plated. Media was replaced every 2 days until cells reached 100% confluency before differentiation. Cells were differentiated by adding the differentiation cocktail (5 μg/mL insulin, 1 μM dexamethasone, 0.5 mM IBMX, 60 μM indomethacin, 1 μM Rosiglitazone). After 48 hours, the media was changed to only include 1 μM Rosiglitazone and 5μg/mL insulin. After another 48 hours, the media was changed to include 5 μg/mL insulin only. On day 5 post differentiation, the cells are considered fully differentiated.

### Treatment of mouse primary white pre-adipocytes with CriPs

***Gfp* target gene.** Primary GFP white pre-adipocytes were isolated from GFP transgenic heterozygous mice and plated in 12 well plates with 8×10^4^ cells per well overnight. Cells were treated with CriPs loaded with *Gfp* sgRNA or control CriPs loaded with Control sgRNA. After 24 hours, cells were fed with fresh culture media. On day 5 post treatment, flow cytometry analysis was performed to measure the loss of GFP. 7-AAD staining was used to determine live cells and dead cells. The percentages of GFP negative cells and GFP positive cells were calculated from the live cells analyzed by flow cytometry. Indels in the *Gfp* genomic locus were measured by T7E1 assay.

***Nrip1* target gene.** Primary white pre-adipocytes were isolated from WT mice and plated in 12 well plates with 8×10^4^ cells per well overnight. Cells were treated with CriPs loaded with each of four different sgRNAs targeting *Nrip1* (*Nrip1* sgRNA 1, *Nrip1* sgRNA 2, *Nrip1* sgRNA 3, *Nrip1* sgRNA 4) or control groups (CriPs- Control sgRNA, Cas9-EP, EP only and non-treated). Cas9-sgRNA: 100 nM. EP: 25 μM. After 24 hours, cells were fed with fresh culture media. Once the pre-adipocytes reached 100% confluency, they were differentiated to adipocytes with the differentiation cocktail. On day 8 post differentiation, cells were collected to measure indels in the *Nrip1* genomic locus by T7E1 assay.

### Gene expression of white adipocytes after *Nrip1* CriP deletion

Primary white pre-adipocytes were isolated from WT mice and treated with CriPs loaded with each of the four sgRNAs targeting *Nrip1* (*Nrip1* sgRNA 1, *Nrip1* sgRNA 2, *Nrip1* sgRNA 3, *Nrip1* sgRNA 4) or control groups (CriPs- Control sgRNA, Cas9-EP, EP only and non-treated). Cas9-sgRNA: 100 nM. EP: 25 μM. Once the pre-adipocytes reached 100% confluency, they were differentiated to adipocytes with the differentiation cocktail. On day 8 post differentiation, cells were collected to measure the expression of UCP1 by RT-PCR. The expression of other thermogenic genes, inflammatory genes and neurotropic factors were also measured in cells treated with CriPs loaded with *Nrip1* sgRNA 3 or Control sgRNA by RT-PCR.

### Off-target effects of CriPs targeting *Nrip1*

Primary white pre-adipocytes were treated with CriPs (Cas9-sgRNA: 100 nM. EP: 25 μM) loaded with *Nrip1* sgRNA 3. Pre-adipocytes were then differentiated to mature white adipocytes. On day 8 post differentiation, cells were collected to measure indels in the *Nrip1* genomic locus or at the off-target sites by T7E1 assay. Sequences targeting the top off-target candidate sites were determined by the CHOPCHOP program(60, 61). Expected DNA bands cleaved by T7E1: On-target: Uncut: 420bp, Cut: 270bp+150bp; Off-target 1: Uncut: 386bp, Cut: 283bp+103bp; Off-target 2: Uncut: 387bp, Cut: 229bp+158bp; Off-target 3: Uncut: 352bp, Cut: 182bp+170bp.

### Cytotoxicity assay post CriPs treatment *in vitro*

The cytotoxicity of Cas9-sgRNA coated with different EP concentrations in different cell types (J774A.1 cells, PECs and primary white pre-adipocytes) was examined by the Vybrant MTT cell proliferation assay. Cells were treated with Cas9-sgRNA (100 nM) coated with different concentrations of EP. After 24 hours, the particle-containing media was replaced by fresh culture media. After another 24 hours, the media was changed to fresh culture medium without phenol red, and 10 μl of 12 mM MTT stock solution was added to each well. The plate was incubated for an additional 4 hours at 37 °C in a humidified CO2 incubator. Following the 4-h incubation, 100 μl of the SDS-HCl solution was added to each well and incubated overnight. The absorbance of the colored formazan product was recorded at 570 nm using a microplate reader (Tecan Group Ltd.) and normalized to the control group with no treatment. An average of three determinations was made.

### *In vivo* treatment of GFP transgenic mice with CriPs

GFP transgenic heterozygous mice (male, 3-8 weeks old) were i.p. injected daily for 5 days with CriPs loaded with *Gfp* sgRNA or Control sgRNA. The CriPs contained 0.9 nmol of Cas9 protein, 0.9 nmol of sgRNA and 20 nmol of EP for each injection. On day 6, mice were sacrificed and the peritoneal cavity was washed with 5 mL of ice-cold PBS to isolate PECs. The cells were plated in the media (DMEM supplemented with 10% (v/v) FBS, 100 μg/mL streptomycin and 100 units/mL penicillin) to enrich for macrophages. Fresh media was added every 48 hours. On day 13 (8 days post last injection), adhered cells were collected. Flow cytometry analysis was performed to measure the loss of GFP. 7-AAD staining was used to determine live cells and dead cells. The percentages of GFP negative cells and GFP positive cells were calculated from the live cells. Genomic DNAs were also collected for detecting indels by deep sequencing.

### Deep sequencing of PECs after *in vivo* CriPs treatment

PECs were isolated from GFP mice i.p. injected with CriPs- *Gfp* sgRNA and CriPs- Control sgRNA. The *Gfp* genomic region of the CriPs- *Gfp* sgRNA target sequence was amplified by PCR using Platinum™ Taq DNA Polymerase High Fidelity kit according to the manufacturer’s protocol. The amplicons were purified using the QIAquick PCR Purification Kit. Libraries were made from the purified amplicons and sequenced on the Illumina MiSeq instrument (300 bp paired-end) by the University of Massachusetts Medical School Deep Sequencing Core Facility. Reads were mapped to the *Gfp* reference sequence, and insertion/deletion/SNP mutations were determined by the CRISPR-Dav program(62).

### Plasma cytokine levels in mice treated with CriPs *in vivo*

WT mice (male, 20 weeks old) were i.p. injected daily for 5 days with CriPs loaded with a non-targeting sgRNA (*Gfp* sgRNA) or PBS. The CriPs contained 0.9 nmol of Cas9 protein, 0.9 nmol of sgRNA and 20 nmol of EP for each injection. Serum was collected at three time points- before injection, 24 hours after the first injection and 2 weeks after the first injection. Plasma cytokine levels (IL-1β, IL-4, IL-6, IL-10, IFNγ, TNFα) were measured using Luminex multiplex assays on a Lincoplex instrument by the National Mouse Metabolic Phenotyping Center at the University of Massachusetts Medical School. The assays were performed according to manufacturer’s recommended procedures. Plasma cytokine levels (IL-10 and TNFα) were also measured in mice treated with LPS for 1.5 hours or without LPS to serve as positive controls for the assay. Data are means ± sem (n = 3-4 mice per group).

### T7E1 assay

Cells were lysed in cell lysis buffer (1M KCl, 1M MgCl2, 1M Tris-Base pH 8.3, 0.45% NP40, 0.45% Tween20, 0.1 mg/ml proteinase K) and then used as templates in PCR reactions to amplify the targeted genomic loci using Platinum™ Taq DNA Polymerase High Fidelity kit according to the manufacturer’s protocol. PCR products were purified using QIAquick PCR Purification Kit and quantified by Nanodrop. Purified PCR products (200 ng) were mixed with 2 μl of 10x NEBuffer 2 (New England Biolabs, Inc) up to a total volume of 19 μl, and denatured then re-annealed with thermocycling at 95 °C for 5 minutes, 95 to 85 °C at 2 °C /second, 85 to 20 °C at 0.2 °C /second. The re-annealed DNA was incubated with 1 μl of T7E1 at 37 °C for 15 minutes. The reaction was stopped by adding 1.5 μl of 0.25 M EDTA, and analyzed on a 4-20% Mini-Protean TBE Gel electrophoresed for 1.5 hours at 100 V, then stained with ethidium bromide. The frequency of indels was calculated based on the band intensities quantified using Image Lab (Bio-Rad Laboratories). The intensities of the cleaved bands were divided by the total intensities of all bands (uncleaved + cleaved) to determine the frequency of indels to estimate gene modification levels.

## ACKNOWLEDGMENT

We thank Joseph Virbasius and members of our laboratory group for excellent discussion of the data in this paper. We appreciate the help of the staff of the Flow Cytometry Core Facility, the Deep Sequencing Core Facility and the National Mouse Metabolic Phenotyping Center (MMPC) at the University of Massachusetts Medical School. We thank Dr. Gang Han for use of the Malvern Zetasizer Nano-ZS particle size analyzer. We also thank Dr. H. Yang for GFP-expressing J774A.1 cells. These studies were supported by grants to M.P.C. from the NIH (DK103047, AI046629, and DK030898) and the International Research Alliance of the Novo Nordisk Foundation Center for Metabolic Research. The National MMPC at UMass is supported by an NIH grant to J.K.K. and M.P.C. (5U2C-DK093000).

## CONFLICT OF INTEREST

The authors declare that they have no conflicts of interest with the contents of this article. The content is solely the responsibility of the authors and does not necessarily represent the official views of the National Institutes of Health.

## AUTHOR CONTRIBUTIONS

Y.S. designed the study, performed the research, analyzed the data and wrote the manuscript. M.P.C designed the study and wrote the manuscript. J.L.C. performed the experiments and analyzed the data. S.M.N. performed the experiments, analyzed the data and revised the manuscript. M.K., B.Y., F.H., and E.T. performed the experiments. Y.J.K.E. analyzed the deep sequencing data. X.H., R.F. and J.K.K. performed the Luminex multiplex assays. All authors reviewed and edited the manuscript.

